# The predictive value of quantitative nucleic acid amplification detection of *Clostridium difficile* toxin gene for faecal sample toxin status and patient outcome

**DOI:** 10.1101/436451

**Authors:** Kerrie A Davies, Tim Planche, Mark H Wilcox

**Author notes:** Corresponding author: Kerrie Davies, Principal Clinical Scientist, Healthcare Associated Infections Research group, Old Medical School, Leeds, LS1 3EX, UK.

## Abstract

**Background:** Laboratory diagnosis of *Clostridium difficile* infection (CDI) remains unsettled, despite updated guidelines. We investigated the potential utility of quantitative data from a nucleic acid amplification test (NAAT) for *C. difficile* toxin gene (tg) for patient management.

**Methods:** Using data from the largest ever *C. difficile* diagnostic study (8853 diarrhoeal samples from 7335 patients), we determined the predicative value of C. difficile tgNAAT (Cepheid Xpert C.diff) low cycle threshold (CT) value for patient toxin positive status, CDI severity, mortality and CDI recurrence. Reference methods for CDI diagnosis were cytotoxicity assay (CTA) and cytotoxigenic culture (CTC).

**Results:** Of 1281 tgNAAT positive faecal samples, 713 and 917 were CTA and CTC positive, respectively. The median tgNAAT CT for patients who died was 25.5 vs 27.5 for survivors (p = 0.021); for toxin-positivity was 24.9 vs 31.6 for toxin-negative samples (p<0.001) and for patients with a recurrence episode was 25.6 vs 27.3 for those who did not have a recurrent episode (p = 0.111). Following optimal cut-off determination, low CT was defined as ≤25 and was significantly associated with a toxin-positive result (P<0.001, positive predictive value 83.9%), presence of PCR-ribotype 027 (P=0.025), and mortality (P=0.032). Recurrence was not associated with low CT (p 0.111).

**Conclusions:** Low tgNAAT CT could indicate CTA positive patients, have more severe infection, increased risk of mortality and possibly recurrence. Although, the limited specificity of tgNAAT means it cannot be used as a standalone test, it could augment a more timely diagnosis, and optimise management of these at-risk patients.

## Background

Laboratory diagnosis of *Clostridium. difficile* infection (CDI) is contentious, especially regarding the use of standalone *C. difficile* toxin gene (tg) nucleic acid amplification tests (tgNAATs) [1–2]. Tg detection is highly sensitive but poorly specific for the diagnosis of CDI. Detection of free toxin within the stool of symptomatic patients, as opposed to toxin gene, correlates with mortality and severity markers [3]. Notably, outcomes of patients who are tgNAAT+ve/toxin-ve are indistinguishable from those who are tgNAAT-ve/toxin-ve [3–4]. The possible over-diagnosis of CDI associated with use of tgNAATs alone has been highlighted, as asymptomatic carriage of *C. difficile* can occurs in up to 20% of inpatients [4]. There is a high prevalence of hospital onset diarrhoea (HOD) from other causes [5], and so without a direct toxin assay, a positive tgNAAT assay may simply reflect the detection of “bystander” organism in a patient colonised by *C. difficile*. Over-diagnosis of CDI in hospitals using standalone tgNAAT can lead to unnecessary treatment, overburden on limited isolation facilities and may cause clinicians to overlook the real underlying diagnosis.

European guidelines recommend *C. difficile* tgNAAT should only be used as part of a two-stage algorithm, with a toxin detection assay as the second stage [2]. Two-stage algorithms using a glutamate dehydrogenase enzyme immunoassay (EIA) or a tgNAAT followed by toxin detection have been widely adopted in the UK [6]. In 2013, 48% of hospitals in 20 European countries were using an optimised algorithm for laboratory diagnosis of CDI, a relative increase of 50% from the previous year [6]. There are distinct differences in laboratory practice between Europe and the USA; in 2014, ^~^44% of US hospitals reported using tgNAAT alone for CDI diagnosis, whilst in Europe (in 2013) that level was much lower at 5% [4,6].

While qualitative results of tgNAATs have poor positive predictive value for CDI, the value of quantitative data from these assays for CDI diagnosis is uncertain. Lower cycle threshold (CT) value (the point at when sufficient nucleotide is detected and the assay is called positive), in molecular assays may indicate greater quantity of assay target; thus, a lower tgNAAT CT value could equate to higher bacterial load. The *C. difficile* tgNAAT CT could provide more information than the binary positive/negative result, by lower CTs predicting poorer outcomes [8]. However, limitations of previous studies (single centre design, lack of optimal reference method, cohort sizes and use of composite outcome) mean that the relationship between CT value and CDI severity, including mortality, and risk of recurrence merits further investigation. We have utilised data from the largest ever study of *C. difficile* laboratory diagnostic assays across four centres to determine the value of the tgNAAT CT value in predicting severity of disease, mortality and CDI recurrence.

## Methods

### Study design

Diarrhoeal faecal samples were collected at 4 UK sites (Leeds Teaching Hospitals, Leeds; St George’s University of London, London; University College Hospital NHS Foundation Trust, London and Oxford University Hospitals NHS Trust, Oxford) between October 2010 and September 2011. Fresh samples were tested with two reference methods: cytotoxicity assay (CTA) and cytotoxigenic culture (CTC), and one commercial molecular method for toxin gene detection; Cepheid Xpert C. diff (Cepheid, Sunnyvale, CA, US) (tgNAAT)) [3]. Patient data were collected for markers of CDI severity (WCC, serum creatinine, serum albumin, and PCR-ribotype) and for patient outcomes (30-day mortality, recurrence and length of hospital stay (LOS)). The molecular assay was performed as per manufacturers’ instructions and results collected for presence of genes for toxin B and binary toxin. PCR-ribotyping was performed as previously described [9]. The study was approved by the National Research Ethics Service (reference number 10/H0715/34).

### Analysis

Only those patients who had at least one NAAT positive sample were included in the analysis. The area under receiver operator curve (AUROC) was calculated for those samples that were positive by tgNAAT to determine if there was an association between low cycle threshold (CT) and patient faecal toxin positive status, CDI recurrence and mortality. Recurrence was defined as a sample that tested positive for toxin (by CTA) at least 14 days after the initial CTA positive sample for that same patient was tested. Recurrence was only assessed in patients who had repeat samples submitted within the study period. Once a low CT had been defined, associations between low CT (≤25) and markers of CDI severity and outcome were investigated by univariate analysis; T-test was used to compare means, Mann-Whitney for Medians (LOS) and Chi-squared for categorical variables.

## Results

There were 8853 samples from 7335 patients, 1281 of which were NAAT positive and could be used for the analysis. Of the 1281, 713 were CTA positive and 917 CTC positive, with a positive agreement of 51.2% (Figure 1). The median tgNAAT CT value for patients who died was 25.5 vs 27.5 for those who survived (p = 0.021), 24.9 vs 31.6 for CTA positive vs CTA negative faecal samples, respectively (p <0.001) and 25.6 vs 27.3 for patients with vs without a laboratory defined CDI recurrence (p = 0.111). CT values of ≤24 (to optimise for CTA positivity) and ≤25 (to optimise for mortality) were both investigated as cut-off values.

**Figure 1.**
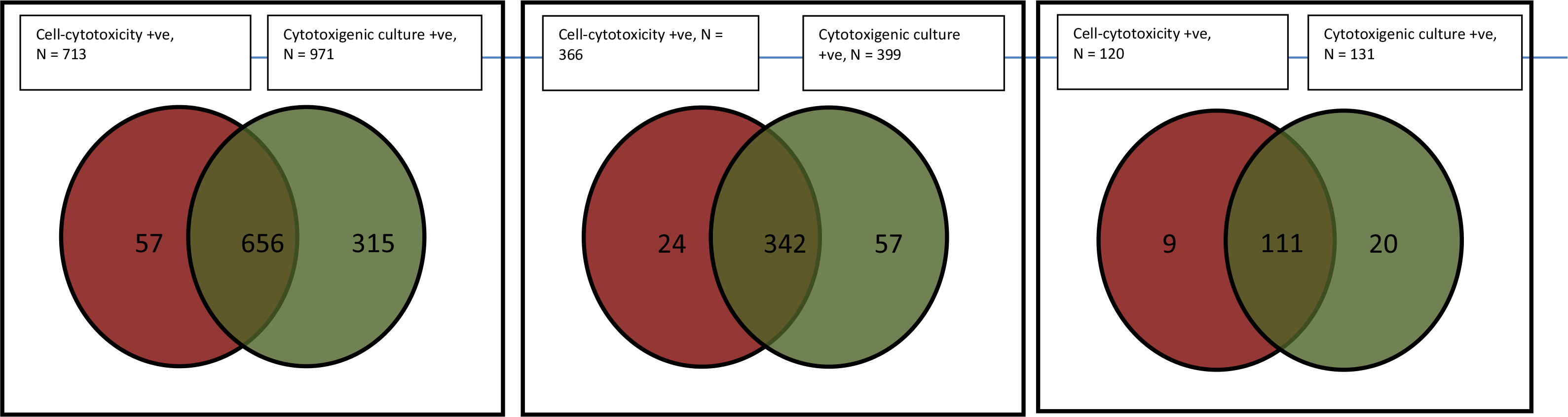
Results of *C. difficile* reference methods A) all tgNAAT +ve samples (n = 1281) positive agreement 51.2%; (B) those with only CT<25 (n = 436) positive agreement 78.4%; (C) those with only CT≤24 (n= 145) positive agreement 76.6%

Of the 436 samples with a tgNAAT CT ≤25, 366 were CTA positive and 399 CTC positive, with a positive agreement of 78.4%. Of the 145 samples with a tgNAAT CT ≤24, 120 were CTA positive and 131 CTC positive, with a positive agreement of 76.6% (Figure 1). The positive predictive values (PPVs) of CT ≤25 and CT ≤24 for recurrence were 49.3% and 18.3%, respectively. The AUROC for CT value and death was 0.572 (95% CI 0.519-0.624, p = 0.009) (Figure 2a); the AUROC for CT value and toxin positive status was 0.831 (95% CI 0.808-0.853, P<0.001) (Figure 2b); and the AUROC for CT value and recurrence was 0.557 (95% CI 0.490-0.624 p = 0.11) (Figure 2c).

**Figure 2.**
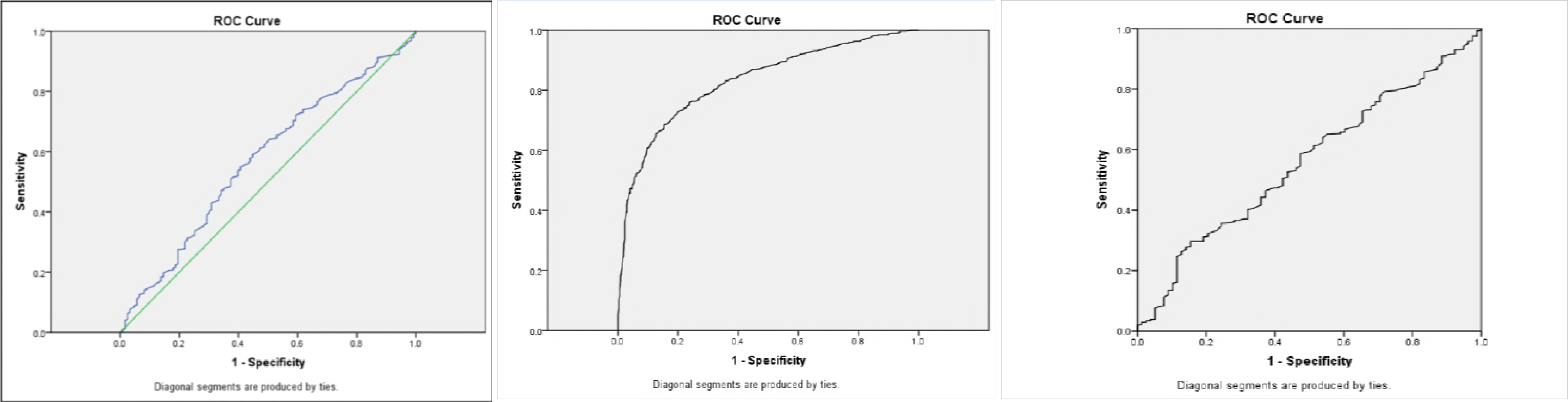
ROCs for CT value against a) mortality (AUROC 0.569) and b) CTA (0.83) and c) recurrence (0.557)

### Low CT

In a univariate analysis, a CT ≤25 was significantly associated with a toxin positive result, presence of PCR-ribotype 027, and mortality with a positive predictive value of 83.9%) for toxin detection (Table 1 and Figure 3). However the sensitivity and specificity for CT≤25 to determine the presence of toxin in tgNAAT positive samples were 51.3% (366/713,95%CI 48-55%) and 87.5% (491/561, 95%CI 84-90%) respectively.

**Table 1.**
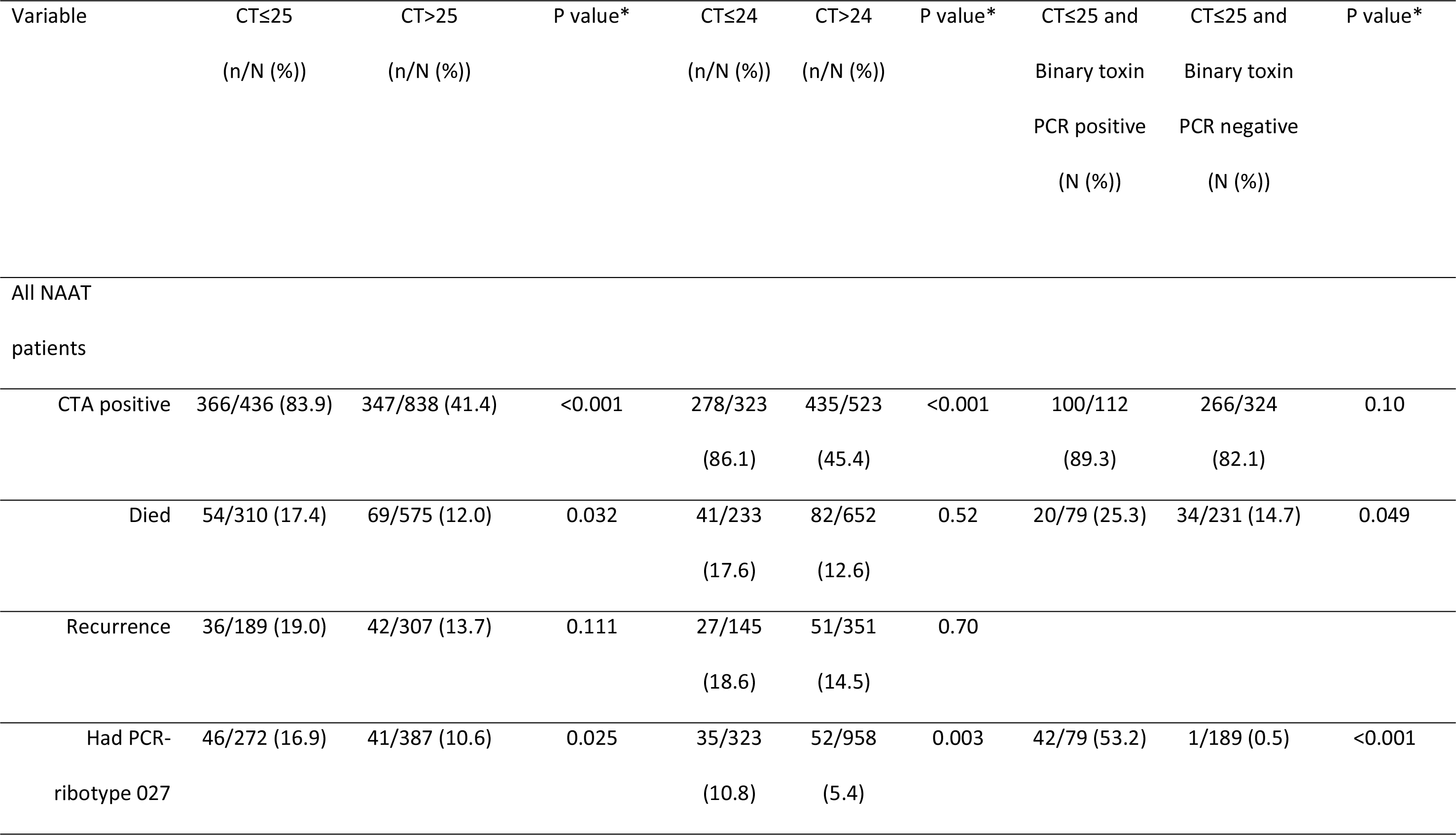
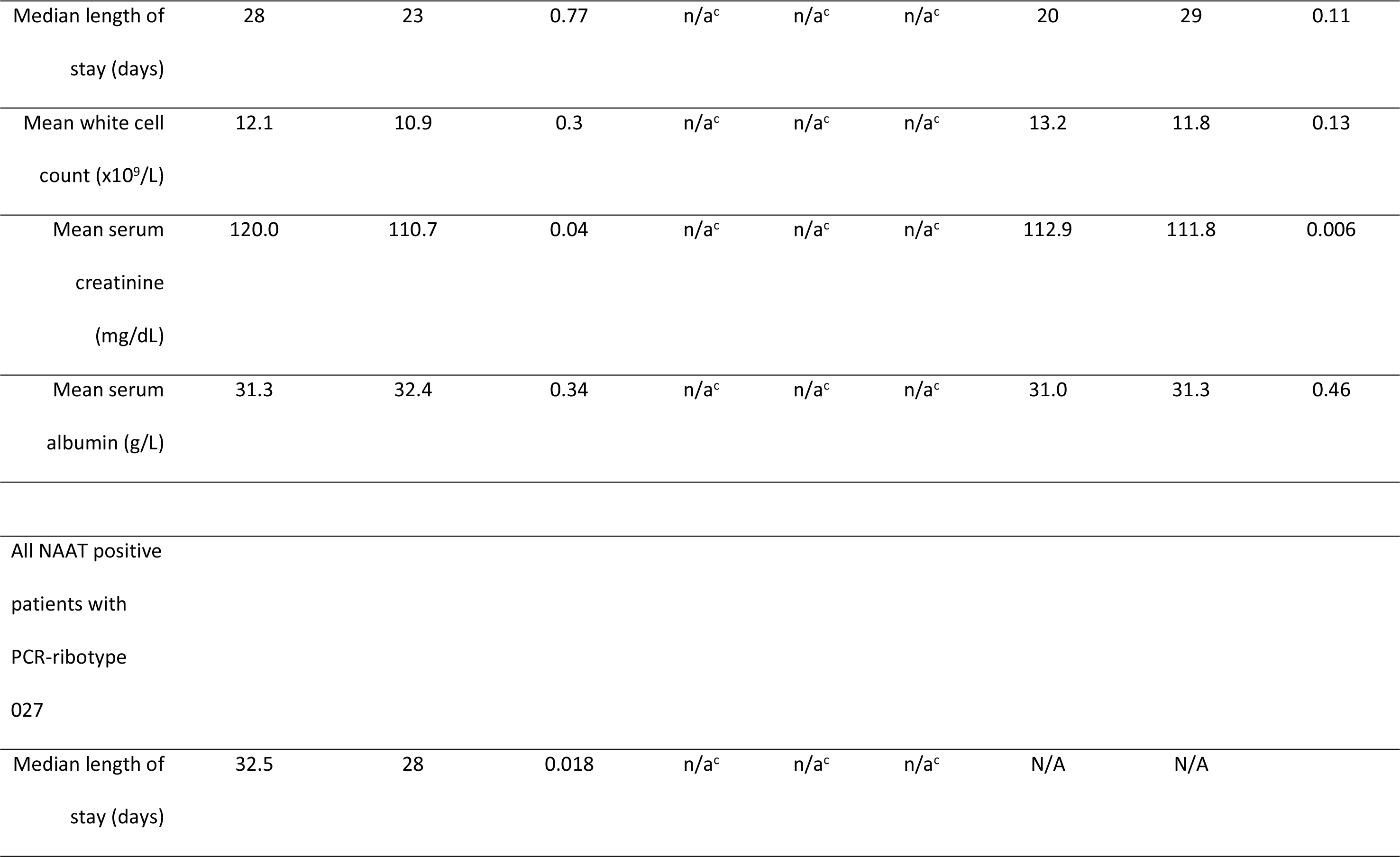
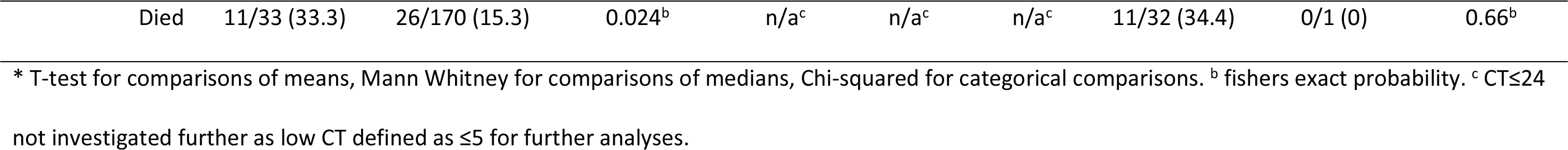
Outcomes and severity markers for patients with CT values ≤25 or > 25

**Figure 3.**
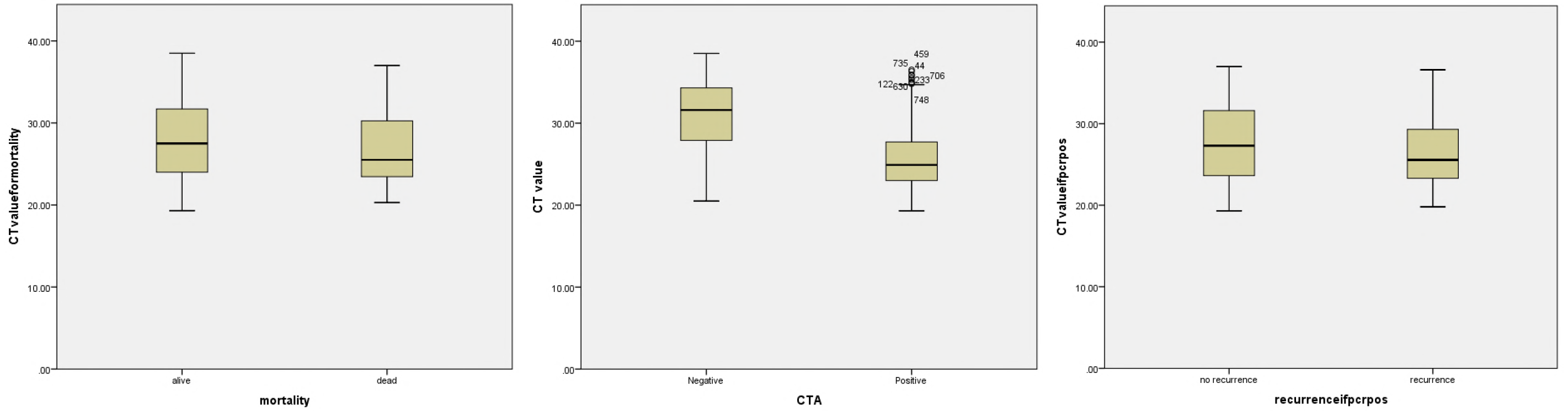
Boxplot of tgNAAT CT value against a) mortality b) CTA result and c) recurrence

A CT ≤24 was also significantly associated with a toxin positive result and the presence of PCR-ribotype 027, with a PPV of 86.1% for toxin detection (Table 1 and Figure 3). Due to the slight advantage of using CT ≤25 over ≤24 as a cut-off, further analyses defined low CT as ≤25.

Patients with a low CT (≤25) had higher mean WCC, a higher baseline mean serum creatinine, lower mean serum albumin and longer length of stay, but these differences were not significant (except serum creatinine p = 0.04, Table 1). For patients with both a low CT and the presence of PCR-ribotype 027, mortality risk and LOS were significantly increased (Table 1). The PPV of low CT for mortality in this population was 17.4% (95% CI 13.5-22.2%); the negative predictive value (NPV) of CT >25 for mortality was 88.0% (65% CI 85.0-90.5%). The relative risk of mortality in CDI with low CT was 1.45 (95% CI 1.0-2.0, p = 0.04), which increased to 2.18 (95% CI 1.2-4.0p = 0.03) in cases also due to PCR-ribotype 027.

The relative risk of mortality in CDI with low CT and a positive binary toxin gene test was 1.72 (95% CI 1.1-2.8, p =0.05), with a significant difference in mortality rates between those with versus without binary toxin gene (25.3 vs 14.7%, p = 0.049, Table 1). There was no difference in the rate of toxin positive (CTA) samples between those with or without detection of the binary toxin gene (Table 1). Patients with a low CT and who were binary toxin positive had higher mean WCC, a higher mean serum creatinine lower mean serum albumin and shorter LOS; of these differences, only increased serum creatinine was significant (p = 0.006) (Table 1). Unsurprisingly, a positive binary toxin gene result was significantly associated with samples containing PCR-ribotype 027 (p<0.001).

### Presumptive 027 result

A PCR-ribotype was available for 817 NAAT positive samples; the sensitivity, specificity, PPV and NPV in this cohort for the Cepheid ‘presumptive 027 ribotype’ result were 93.5 % (95% CI 85.8-97.3), 94.6% (95% CI 92.6-96.1), 68.8% (95% CI 59.8-76.6) and 99.1% (95% CI 98.0-99.6), respectively. Those samples that were falsely identified as containing ribotype 027 did not contain strains with ribotypes that fall within the same clade as 027 (data not shown), and so are not known to have the same tcdC deletion as used to presumptively identify 027 in this assay.

## Discussion

This is the largest study to date examining the clinical utility of tgNAAT CT value to predict the diagnosis and outcomes of CDI. Importantly, this is the only study that has tested all samples with both the CTA and CTC reference assays. We have shown that a low tgNAAT CT value (≤25) is significantly associated with a toxin positive status, presence of PCR-ribotype 027, and mortality. Laboratory defined CDI recurrence was also associated with low CT, although this was not significant, possibly due to the low number of such cases within the dataset. The median tgNAAT CT value for patients who died was significantly lower than for those who survived (25.5 vs 27.5 p = 0.021) with a significant AUROCC for NAAT positives and mortality; 0.572 p =0.009. In addition, the median tgNAAT CT value for a diarrhoeal sample that was (CTA) toxin positive was 24.9 compared with 31.6 for toxin negative samples (p <0.001); the AUROC here was 0.83 (p <0.001).

In order to optimise the CT cut-off we investigated both ≤24 and ≤25 cycles but found little difference between these (Table 1); a slighter higher PPV for toxin positivity using CT ≤24 was offset by the lack of significant association with mortality seen using CT threshold ≤25. We therefore used CT ≤25 as our ‘low CT’ threshold, which was slightly higher than found in two smaller recent studies [8,10]. One of these reported an AUROC of 0.857 when using a CT of 23.5; however, this was for a composite end-point of ‘poor outcome’ [8,]. As our patient cohort was ^~^8-fold larger than in the previous study, we have been able to determine a significant association between low CT and single outcome measures, rather than for a composite outcome. Indeed, as the PPV of a low tgNAAT CT (≤25) for toxin detection was 83.9%, it is possible that such a threshold could be used as a proxy for presumptive toxin detection, as suggested from a recent small study [11]. However, it is difficult to recommend tgNAAT be used as a standalone test, even with these cut-offs, as it has relatively low sensitivity and specificity for confirmation of detection of toxin gene (51% and 88%) and only moderate usefulness in the prediction of mortality (AUCROCC 0.568 and relative risk of death 1.45). Qualitative (positive/negative) and higher quantitative CT values in this tgNAAT are even less predictive of toxin positivity status [3,4].

The relative risk of mortality in patients with a low CT (≤25) vs a CT>25 was 1.45 which increased to 2.18 in those patients who had a low CT and had PCR-ribotype 027. Indeed, in those patients with a low CT (≤25), mortality and longer LOS were also significantly associated with the presence of PCR-ribotype 027, compared with other PCR-ribotypes, highlighting a possible ‘at risk’ group. There is conflicting evidence regarding the role of binary toxin in CDI outcomes [12–13]. Our data show that the relative risk for mortality in patients with low CT increased from 1.45 to 1.72 when the presence of binary toxin gene was taken into account. Patients with a low CT also had a lower serum albumin but mean WCC, higher mean serum creatinine, longer LOS were not significantly different. This was possibly due to the small sample size in these subgroups. The same pattern was seen in patients with a low CT who were also binary toxin gene positive, with the exception that these had a shorter LOS than binary toxin negative subjects. This warrants further investigation in a larger, prospective study.

Prevention of recurrence is a key goal of patient management in cases of CDI, indeed the antibiotic fidaxomicin is recommend in first recurrent cases to prevent further recurrence [14], due to the reduction in recurrence risk with this antibiotic compared with vancomycin [15]. The high acquisition cost of some CDI treatment options means that ways are needed to predict patients who have higher risk of recurrence or other poor outcomes. Although CDI recurrence was associated with a low CT in our study, this was not a significant finding, possibly due to the low number of recurrent cases within this dataset. Risk of CDI recurrence has also previously been shown to be higher in PCR-ribotype 027 [16[. However, traditional typing methods used to identify the ribotype of a strain causing CDI require culture first and so are not timely; in this regard, a rapid ‘presumptive 027’ result could be of value. We found that the Cepheid Xpert C. diff assay has a high NPV for ‘presumptive RT027’, but does overcall the number of samples that truly contain PCR-ribotype 027 (i.e. a low PPV of 69.9%). The presence of the gene target used to identify PCR-ribotype 027 by the Cepheid Xpert C. diff assay in other ribotypes related to PCR-ribotype 027 can confound the results produced by this assay [17–18].

Our study has some limitations. We performed a retrospective data analysis and had no validation set to test. Nevertheless, we utilised data from the largest ever study of *C. difficile* diagnostic assays [3], which contemporaneously collected CDI outcome data. Our analyses of recurrence were based on a laboratory definition (i.e. submission of more than one patient sample), which will likely miss true cases; for example, patients who did not have a new sample submission, or who were retested elsewhere [3]. Our original study was not designed to capture recurrence data systematically and a full prospective study of the impact of CT value information on recurrence rates and patient management is warranted. Additionally, our data only relate to one NAAT, and so require confirmation with respect to other assays. However, there is some emerging evidence that other NAATs may have similar quantitative predictive value [19].

Developing predictive tools is a step forward, but also a challenge to patient treatment. Our study provides evidence of how quantitative tgNAAT results could augment the diagnosis and management of CDI. Low tgNAAT CT values (≤25) indicate patients with more severe infection and at higher risk of mortality and possibly recurrence. In those with a low CT (≤25) and a presumptive RT027 result, the risk of mortality increases further. Timely diagnosis and optimised management may have the biggest impact in these patients.

## Acknowledgments

We thank all of the sites and staff that took part in the original study; Leeds Teaching Hospitals, Leeds; St George’s University of London, London; University College Hospital NHS Foundation Trust, London and Oxford University Hospitals NHS Trust, Oxford.

